# PrP^Sc^-induced conformational changes and strain-specific structures of PrP^Sc^ revealed by Disulfide-crosslink scanning

**DOI:** 10.1101/175919

**Authors:** Yuzuru Taguchi, Noriyuki Nishida, Hermann S. Schatzl

**Affiliations:** Department of Comparative Biology & Experimental Medicine, Faculty of Veterinary Medicine, University of Calgary, Calgary, Alberta, Canada; Division of Cellular and Molecular Biology, Department of Molecular Microbiology and Immunology, Nagasaki University Graduate School of Biomedical Sciences, Nagasaki, JAPAN; Departments of Molecular Biology and of Veterinary Sciences, University of Wyoming, Laramie, Wyoming, U.S.A.

**Keywords:** prion protein, prion conversion, protein cross-linking, protein misfolding, protein structures

## Abstract

There exist many phenotypically-varied prion strains, like viruses, despite the absence of conventional genetic material which codes the phenotypic information. As prion is composed solely of the pathological isoform (PrP^Sc^) of prion protein (PrP), the strain-specific traits are hypothesized to be enciphered in the structural details of PrP^Sc^. Identification of the structures of PrP^Sc^ is therefore vital for the understanding of prion biology, though they remain unidentified due to the incompatibility of PrP^Sc^ with conventional high-resolution structural analyses. Based on our previous hypothesis that the region between the first and the second α-helix (H1∼H2) and the distal region of the third helix (Ctrm) of the cellular isoform of PrP (PrP^C^) have important roles for efficient interactions with PrP^Sc^, we created series of mutant PrPs with two cysteine substitutions (C;C-PrP) which were systematically designed to form an intramolecular disulfide crosslink between H1∼H2 and Ctrm and assessed their conformational changes by prions: Specifically, a cysteine substitution in H1∼H2 from 165 to 169 was combined with cysteine-scanning along Ctrm from 220 to 229. C;C-PrPs with the crosslinks were expressed normally with the similar glycosylation patterns and subcellular localization as the wild-type PrP albeit with varied expression levels. Interestingly, some of the C;C-PrPs converted to the protease-resistant isoforms in the N2a cells persistently infected with 22L prion strain, whereas the same mutants did not convert in the cells infected with another prion strain Fukuoka1, indicating that local structures of PrP^Sc^ in these regions vary among prion strains and contribute to prion-strain diversity. Moreover, patterns of the crosslinks of the convertible C;C-PrPs implied drastic changes in positional relations of H1∼H2 and Ctrm in the PrP^Sc^-induced conformational changes by 22L prion. Thus, disulfide-crosslink scanning is a useful approach for investigation of strain-specific structures of PrP^Sc^, and would be applicable to other types of amyloids as well.

---

Prions are pathogens composed solely of aberrantly folded isoforms (PrP^Sc^) of cellular prion protein (PrP^C^) devoid of any nucleotide genome which usually codes pathogenic information. Prions cause fatal neurodegenerative disorders in various mammalian species, e.g. Creutzfeldt-Jakob disease (CJD) in humans, scrapie in sheep and goat, chronic wasting disease (CWD) in cervids and bovine spongiform encephalopathy in cattle [1]. Despite the lack of a nucleotide genome, prions behave like viruses in terms of *quasi-species* nature, high specificity of host ranges and diversity in clinicopathological features which are stably inherited over generations [2]. Their pathogenic characteristics are thought to be enciphered in the structures of PrP^Sc^ [3] and high-fidelity representation of the structures on the nascent PrP^Sc^ through a template-guided refolding of PrP^C^ by the template PrP^Sc^ enables faithful inheritance of the traits. Elucidation of details of the structures of PrP^Sc^ and the refolding process is therefore essential for prion research, but they have not been unveiled yet because PrP^Sc^ is unsuitable for conventional high-resolution structural analyses. Alternatively, structural models of PrP^Sc^ were deduced based on secondary-structural information of PrP^Sc^ obtained with Fourier transform infrared spectroscopy, hydrogen/deuterium exchange analysis [4][5] or images of electron microscopy on two-dimensional crystals or fibrils of purified PrP^Sc^ [6][7][8]. Structural differences between prion strains were also inferable from varied biochemical properties of PrP^Sc^, e.g. molecular size of proteinase K (PK)-resistant fragments (PK-res)[9], structural stabilities in denaturant solutions [10][11] and glycoform ratios [12]. As another approach, Hafner-Bratkovic et al utilized disulfide crosslinking of recombinant PrPs to identify regions which retain their structures through the *in vitro* aggregation formation process [13].

Unlike PrP^Sc^, high water-solubility and small molecular size of PrP^C^ allowed detailed structural analysis by nuclear magnetic resonance spectroscopy (NMR). The global three-dimensional structures of PrP^C^ are highly conserved among different species with the same secondary-structure components, i.e. two short beta strands (**Fig. 1A;** B1 and B2) and three alpha helices (H1, H2 and H3) [14][15][16]. Interspecies variation in amino-acid sequences tend to cluster at some spots including the region between H1 and H2 (H1∼H2) or near the C-terminal glycosylphosphatidylinositol (GPI) anchor-attachment site (Ctrm) [17], which often affect the interspecies transmission of prion [18][19]. For example, an asparagine at the codon 170 can greatly affect inter-species transmissions of prions; transmission of CWD to transgenic mice expressing an elk/mouse chimeric PrP with mouse residues only in Ctrm was substantially inefficient [20]. Moreover, a polymorphism in Ctrm of cervid PrP influences the stability of CWD strains [21].

**FIGURE 1.**
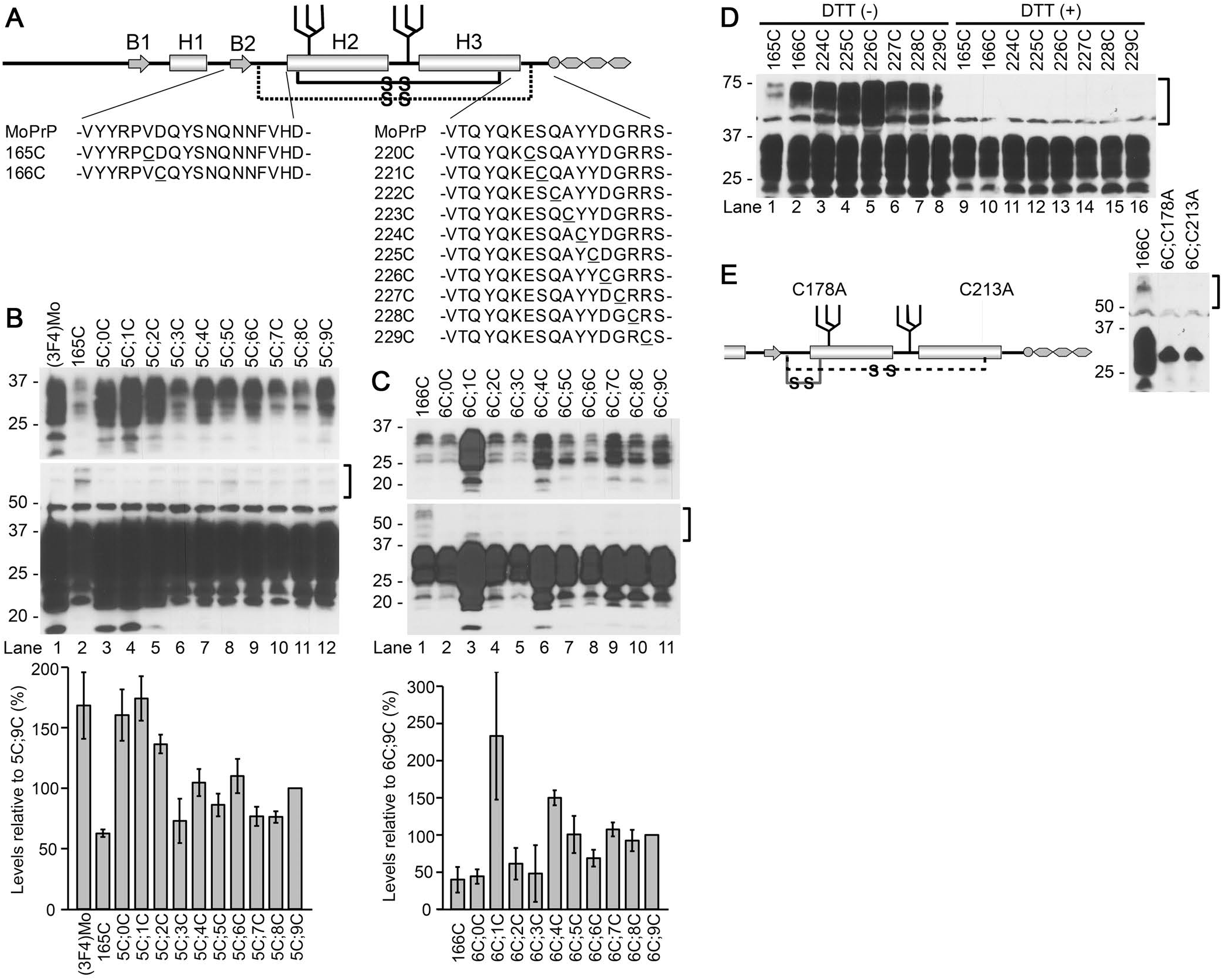
Design and expression of mutant PrPs with two cysteine (Cys) substitutions, 165C;C- and 166C;C-series. **A.** A schematic illustration of the secondary-structure components of mouse PrP and positions of the substituted Cys. MoPrP, sequences of wild-type mouse PrP. B1 and B2, the first and the second β strands, respectively. H1, H2 and H3, the first, second and third a helices[45]. 165C or 166C was combined with another Cys scanning the distal H3 (Ctrm) from 220 to 229 (165C;C- or 166C;C-series, respectively; substituted Cys are underlined). The solid and broken lines with “-S-S-“ represent the native and the newly-introduced disulfide crosslink, respectively. **B and C.** Expression levels and banding patterns of 165C;C- or 166C;C-series. Immunoblots with 3F4 monoclonal antibody (mAb) of whole-cell lysates of transiently-transfected N2a cells. Each two-Cys construct is named after the last digits of the residue numbers of the substituted Cys (in mouse numbering). 165C or 166C, mutants with Cys only at 165 or 166, respectively. Square brackets, positions of dimeric forms of the mutant PrPs. The upper panels and lower panels of blots represent images of short- and long-exposure to the same PVDF membranes, respectively. Note that all the constructs have the similar banding patterns as (3F4)MoPrP, indicating the complex-type N-linked glycosylation. Two-Cys mutants show trace amounts of dimers whereas 165C or 166C have discernible dimers despite their low expression levels. Bottom panels are graphs showing expression levels of each series quantified by densitometry. Each bar represents mean ± standard deviation from three independent experiments. **D** Single-Cys PrPs have substantial levels of dimers which disappear by dithiothreitol (DTT) treatment. Immunoblots with 3F4 mAb showing banding patterns of mutant PrPs with only a single Cys substitution either in H1∼H2 or Ctrm. DTT (+) and (-), samples with or without DTT in the sample buffer. The square bracket indicates the position of the dimeric forms. **E** Disruption of the native disulfide bond drastically changes the banding pattern, suggesting the substituted Cys of 165C;C- or 166C;C-series do not affect the native disulfide bond. **Left panel.** A schematic illustrating positions of alanine substitutions for the native Cys. The solid or broken lines with “-S-S-“ in the schematic illustration represents putative disulfide crosslinks of 6C;C213A or 6C;C178A, respectively. **Right panel.** Immunoblots with 3F4 mAb showing banding patterns of the mutant PrPs combining 166C with the alanine substitution. Note that the banding patterns of 6C;C178A and 6C;C213A are very different from that of 166C, presumably high-mannose-type N-glycosylation, unlike the banding patterns of 165C;C- or 166C;C-series constructs.

We previously demonstrated that efficiencies of dominant-negative inhibition by mutant PrPs which have internal deletions in H1∼H2 correlated with the deletion sizes, propounding that H1∼H2 might be an interaction interface for PrP^C^-PrP^Sc^ interactions [22]. Besides, positional relations between H1∼H2 and Ctrm seemed important for the deletion mutants to efficiently interact with PrP^Sc^. Inspired by those findings, we hypothesized that positional relations of H1∼H2 and Ctrm are influential on PrP^C^-PrP^Sc^ interactions and the subsequent conversion. To test this hypothesis, we created series of mutant PrPs with two cysteine (Cys) substitutions (C;C-PrP), one in H1∼H2 and the other in Ctrm, which crosslink the two regions by an artificial disulfide bond, and evaluated their effects on the PrP^C^-PrP^Sc^ conversion. Those intramolecularly-crosslinked PrPs were normally expressed on the cell surface and, when expressed in N2a cells persistently infected with a mouse-adapted scrapie 22L (22L-ScN2a), some of them converted into PK-res isoforms in a PrP^Sc^-dependent manner. Interestingly, convertibility of the mutants crucially depended on certain patterns of crosslinks. Furthermore, the convertibility of C;C-PrP seemed to be strain-dependent, suggesting that this region is responsible for prion-strain diversity. Our unique approach provides novel insights into the structural requirements for PrP^c^-PrP^Sc^ conversion.

## EXPERIMENTAL PROCEDURES

### Reagents and antibodies

All media and buffers for cell culture and Lipofectamine LTX Plus were from Life Technology Corporation (Carlsbad, CA, USA). Plasmid purification kit, DNA gel extraction kit, Site-directed mutagenesis kit, detergents [including Triton X-100 (TX100), deoxycholic acid (DOC), Triton X-114 (TX114), Tween 20 and sodium dodecyl sulfate (SDS)], proteinase K (PK), anti-PrP monoclonal antibodies (mAb) 4H11 and 3F4 (recognizing residues 108–111 of human PrP), and all secondary antibodies were as previously reported [22]. Iodoacetamide (IAA), dithiothreitol (DTT), Glu-C endopeptidase (V8 protease), and anti-FLAG polyclonal antibody were purchased from Sigma-Aldrich Co., LLC (St. Louis, MO, USA).

### Site-directed mutagenesis

All primers for site-directed mutagenesis were ordered from Integrated DNA Technologies, Inc. (Coralville, IA, USA) and are listed in **Table S1**. Mutations were made with QuickChange Site-Directed Mutagenesis Kit (Agilent Technologies, Inc., Santa Clara, CA, USA) according to the manufacturer’s instruction. Sequences of mutant PrPs were determined by Eton Bioscience, Inc. (San Diego, CA, USA).

### Cell culture, transient transfection and analysis of PK-resistant fragments

Transient transfection of mouse neuroblastoma cell lines with or without persistent scrapie infection (22L-ScN2a or N2a, respectively), and procedures for preparation of samples of transfected N2a or 22L-ScN2a cells were as previously described [22], except for some modifications. Briefly, cells on 24-well plates were transfected with 0.3 μg/well of each plasmid with Lipofectamine LTX Plus (Life Technologies) for evaluation of expression or PK-res levels of mutant PrPs. For evaluation of dominant-negative inhibition, 0.2 μg each of (3F4)MoPrP and mutant PrP were co-transfected. The Fukuoka1- and RML-infected N2a58 cells were also previously described [23][24].

### SDS-polyacrylamide gel electrophoresis (SDS-PAGE) and immunoblotting

The protocol for SDS-PAGE, development of blots, methods of densitometry and quantification have been described previously [22].

### Digestion with V8 protease

N2a cells, ∼60 % confluent on 6-well culture plates, were transiently transfected with 1.0 μg/well of plasmid coding the mutant PrP with Lipofectamine LTX. Next day, the medium was replaced with fresh medium and cells were cultured further at 37°C in a CO_2_ incubator. 48 hours after transfection, cells were rinsed once with phosphate-buffered saline (PBS) and then 1 ml/well of 1.5 mM IAA in PBS was overlaid and incubated for 10 minutes at 4°C. After removal of IAA, cells were rinsed once with PBS without calcium and magnesium (Ca/Mg) and incubated in 700 μl/well of 3mM EDTA in PBS without Ca/Mg at 4°C for 5 minutes. Then, the cells were mechanically detached by pipetting and collected in a tube. The cell suspension was centrifuged at 1,000 x g at 4°C for 5 minutes and the supernatant was discarded. 400 μl of phosphate-buffered 2% Triton X-114 (TX114) lysis buffer was added, cells were resuspended by vortexing for ∼10 seconds, and incubated on ice for 30 minutes, with a few seconds of vortexing from time to time. The lysate was then centrifuged at 16,100 x g at 4°C for 1 minute and the supernatant transferred to a screw-cap tube as TX114 lysate. PrP was concentrated by TX114 extraction and methanol/chloroform precipitation as previously described [22]. The pelleted proteins after methanol/chloroform precipitation were dissolved in 0.5% SDS in 50 mM sodium bicarbonate on a shaking incubator (Thermomixer; Eppendorf AG, Germany), at 95°C for 10 minutes with shaking at 1,400 rpm. After the pellet was completely dissolved, the solution was diluted with a 4-fold volume of 200 mM sodium bicarbonate to dilute SDS concentration, so that V8-protease efficiently digests PrP. After addition of 2 μl of V8 protease (2.5 U/μl), the solution was incubated at 37°C for 1 hour. Finally, 1/4-volume of 5 x sample buffer with or without DTT was added and boiled. For re-probing of the PVDF membrane, the membrane was incubated in 100 % methanol for 20 minutes, washed in TBST and incubated with another primary antibody in 5% milk in TBST.

### Immunofluorescence analysis

The procedures for transient transfection of cells, fixation, permeabilization, and immunolabeling were as reported previously [22], except that samples were analyzed on an epifluorescence microscope, Olympus IX51, with objective lens Olympus LUCPlanFL N 40 x (0.60), and images were acquired with software Olympus DP2-BSW.

## RESULTS

### Design of C;C-PrP series

In order to assess the significance of the intramolecular interactions between H1∼H2 and Ctrm on PrP^C^-PrP^Sc^ interactions and the subsequent conversion, we created series of mutant PrPs which have two Cys substitutions, one in H1∼H2 and the other in Ctrm so that the two regions are cross-linked by an artificial disulfide bond (**Fig. 1A**), and tested their conversion to PK-res forms. For the Cys substitution in H1∼H2, we selected Val165 and Asp166 (residues were numbered according to mouse numbering unless otherwise noted), because they are close enough to Ctrm to form a stable disulfide crosslink in a native PrP^C^ conformation (PDB ID: 2L39) [14]. Since the global conformation of a mutant human PrP with an extra disulfide bond between residues 166 and 221 (in human numbering; they are equivalent to 165 and 220 of mouse PrP, respectively) was indeed similar to that of wild-type human PrP [25][26], we expected that the same holds for mouse PrP. The second Cys substitution scanned Ctrm from the residue 220 to 229. C;C-PrP constructs are named after the positions of Cys but only the last-digit numbers were used for simplicity, e.g. a mutant with Cys at 165 and 229 is named as “5C;9C”. Since a 3F4 epitope-tagged mouse PrP [(3F4)MoPrP] was used as the template for site-directed mutagenesis, every mutant PrP carries a 3F4 epitope tag.

### Expression levels, glycosylation and subcellular localization of C;C-PrP series

We transfected N2a mouse neuroblastoma cells with the plasmids coding 165C;C- and 166C;C-series (**Fig. 1A**) to examine expression levels and glycosylation status of the mutant PrPs. Banding patterns of all mutants were similar to that of (3F4)MoPrP (**Fig. 1B**), typical of PrP^C^ with complex-type N-linked glycans and GPI anchor, and lacked the dimeric forms (**Fig. 1B,** square bracket). Their expression levels were varied (**Fig. 1B,** graphs). A C;C-PrP which has Cys residues equivalent to those of the aforementioned human PrP mutant [25], namely 5C;0C, showed the highest expression level comparable to (3F4) (**Fig. 1B,** lane 2) presumably because its intramolecular disulfide crosslink between the substituted Cys did not interfere with the native PrP^C^ conformation as discussed later. All the 166C;C-series mutants showed similar banding patterns as 165C;C-series without discernible dimeric forms (**Fig. 1C,** square bracket). 6C;1C and 6C;4C showed highest expression levels among 166C;C-series (**Fig. 1C,** lanes 3 and 6). “Intramolecular” disulfide crosslink of C;C-PrPs is implied by the absence of discernible dimeric forms; unlike C;C-PrPs, all the mutant PrPs with a single Cys substitution formed substantial levels of dimeric forms which are presumably crosslinked by an ‘intermolecular’ disulfide bond and disappear upon dithiothreitol (DTT) treatment [**Fig. 1D,** DTT(-) vs (+)].

To rule out the possibility that the Cys residues at 178 and 213 which contribute to the native disulfide bond might be shuffled to couple with the substituted Cys, we replaced either Cys at 178 or 213 with alanine so that the native disulfide bond is broken and instead coupled with 166C (6C;C178A and 6C;C213A) (**Fig. 1E,** schematic). The banding patterns of those mutants were very different from that of wild-type PrP, reminiscent of PrP with high-mannose-type N-linked glycans [27] (**Fig. 1E,** right panel). The absence of those features supported that C;C-PrPs form an intramolecular disulfide crosslink between the substituted Cys without affecting the native disulfide and undergo normal folding and processing in ER and trans-Golgi network. In accordance with the view, immunofluorescence analysis demonstrated that C;C-PrPs were distributed on the cell surface (**Fig. 2,** non-permeabilized) and in the perinuclear region as clusters (**Fig. 2,** permeabilized) like wild-type PrP, corroborating normal intracellular trafficking and subcellular localization of C;C-PrPs.

**FIGURE 2.**
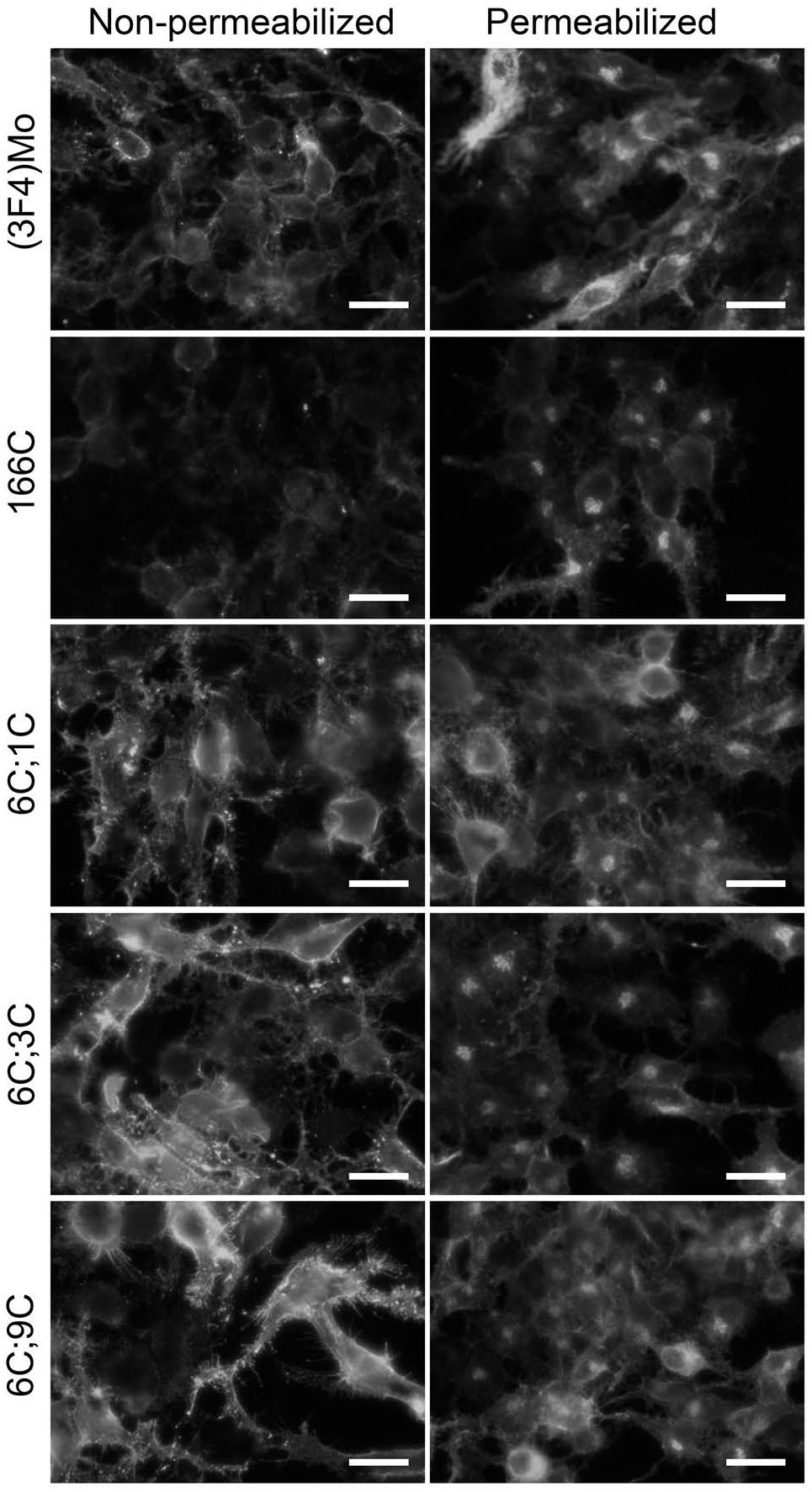
166C;C-series mutants show the same subcellular localization as the wild-type mouse PrP. Epifluorescence microscopic images of immunofluorescence with 3F4 mAb with or without permeabilization. All of the C;C-constructs exhibit the same localization patterns as the wild-type, i.e. on the cell-surface and in the perinuclear regions. Scale bars, 25 μm.

### Evidence for intramolecular disulfide crosslink formation

To demonstrate intramolecular disulfide crosslink formation by the substituted Cys, we introduced a FLAG-tag to C;C-PrPs (**Fig. 3A**) and analyzed the fragment patterns of V8 protease-digested products. A crosslink between H1∼H2 and Ctrm theoretically produces extra bands on immunoblots by bonding fragments (**Fig. 3A**). Indeed, V8-digested FLAG-tagged (3F4)MoPrP, 166C, 6C;3C and 6C;9C (**Fig. 3B**) showed distinct banding patterns along with findings suggestive of intramolecular disulfide crosslink: First, full-length 6C;3C-FLAG and 6C;9C-FLAG remained after the digestion (**Fig. 3B,** compare lanes 7 and 8, arrowhead), whereas full-length (3F4)Mo-FLAG and 166C-FLAG PrP were completely digested (**Fig. 3B,** compare lanes 5 and 6, arrowhead). Relative protease resistance of C;C-PrPs was also implied by smaller amounts of fragments produced by endogenous proteolysis (**Fig. 3B,** lanes 3 and 4, square bracket). These are attributable to steric effects caused by the crosslink of H1∼H2 and Ctrm concealing protease-vulnerable regions. Second, the greatly improved immunoreactivity of the smallest fragments of 6C;3C-FLAG and 6C;9C-FLAG by DTT treatments (**Fig. 3B,** lanes 7 *vs.* 11 or 8 *vs.* 12) also indicates the steric effects hiding the epitope and its re-exposure by DTT which breaks apart the crosslinked fragments. Third, the intermediate size fragments of V8-digested 6C;3C-FLAG and 6C;9C-FLAG (**Fig. 3B,** arrowhead and square bracket, respectively) which disappeared by DTT (**Fig. 3B,** lanes 7 and 8, curled bracket) would apparently represent the predicted “extra fragments” (**Fig. 3B,** lanes 5 and 6, square bracket). Taken together, these findings strongly support the intramolecular crosslink formation between the Cys residues.

**FIGURE 3.**
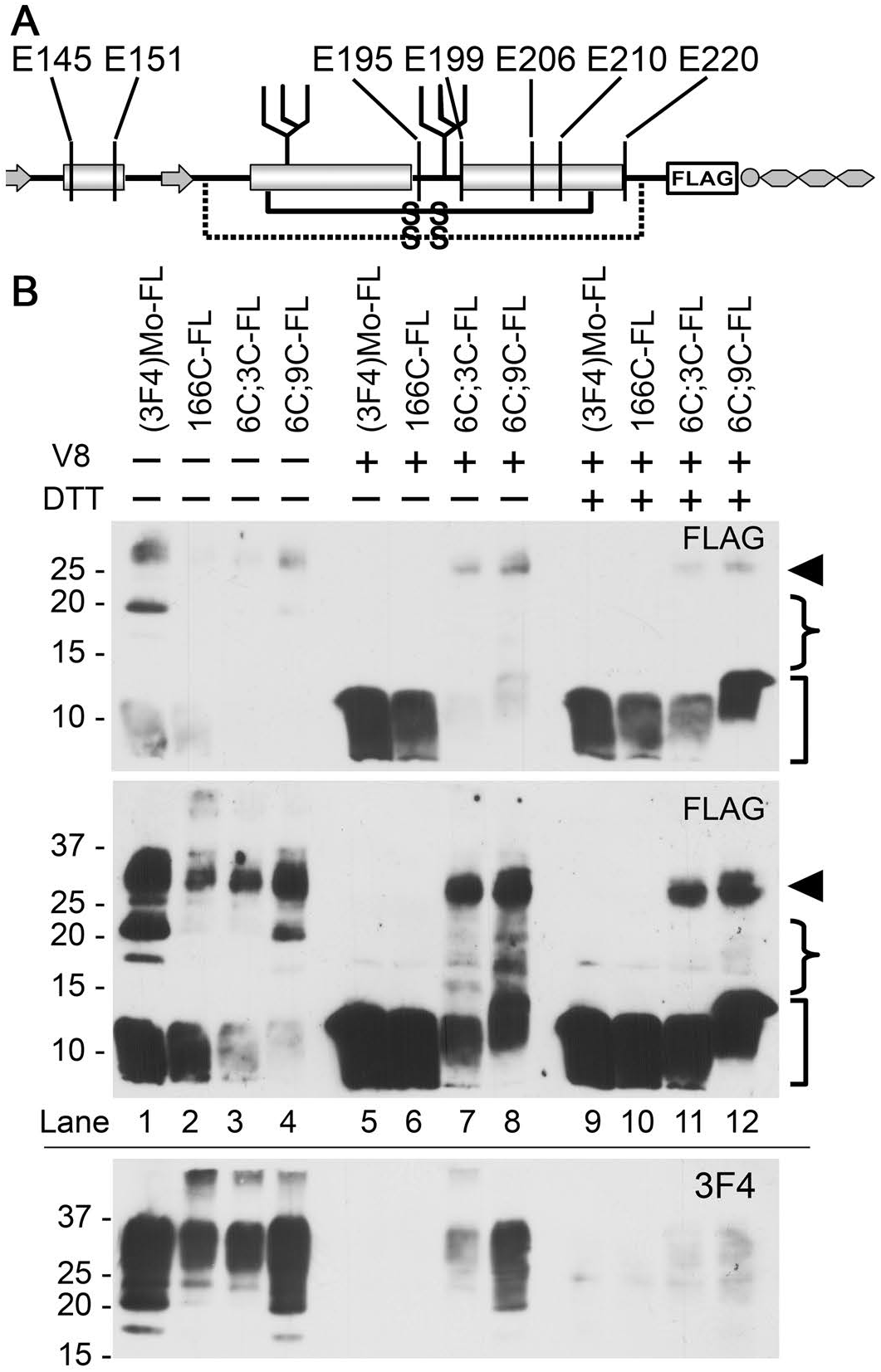
Substituted Cys of 166C;C-constructs form an intramolecular disulfide crosslink: assessment of fragment patterns after V8 protease digestion. **A** schematic illustrating the position of the FLAG tag of the FLAG-tagged 166C;C-constructs along with putative cleavage sites by V8 protease (vertical lines). **B**. V8-digested fragment profiles are similar between the FLAG-tagged wild-type [(3F4)Mo-FL] and 166C-FLAG (166C-FL) but those of 6C;3C-FLAG and 6C;9C-FLAG (6C;3C-FL and 6C;9C-FL, respectively) are different. Immunoblots with anti-FLAG polyclonal antibody, or 3F4-mAb, showing non-digested and V8-digested PrPs with or without DTT in the sample buffer. The upper-panel and middle-panel images were obtained from the same PVDF membrane with shorter and longer exposure, respectively. The bottom panel is immunoblots reprobed with 3F4-mAb. Arrowhead, full-length FLAG-tagged PrP molecules. Curled brackets, positions of the intermediate-size fragments that diminish by DTT. Square brackets, smallest fragments.

### Conversion of C;C-PrPs into PK-res isoforms by bona fide PrP^Sc^

Next, we assessed conversion efficiencies of 165C;C- and 166C;C-series mutants by expressing them in 22L-ScN2a and evaluating their PK-res. Among 165C;C-series, only 5C;8C and 5C;9C showed PK-res (**Fig. 4A**), while 166C;C-series exhibited gradually increasing levels of PK-res from 6C;5C to 6C;9C **(Fig. 4B**). Just as PrP^C^ isoforms, PK-res of C;C-PrPs lacked dimeric forms (**Fig. 4C,** Double-Cys), whereas every single-Cys PrPs tested showed intense dimeric bands (224-229C; **Fig. 4C,** Single-Cys) which disappeared with DTT (**Fig. 4D)**. These data support the view that the PK-res of C;C-PrPs were not derived from PrP^C^ isoform with free Cys residues, i.e. without crosslink, but from those with the intramolecular disulfide crosslink. The absence of PK-res in non-infected N2a cells demonstrated that the conversion of 6C;9C into PK-res isoform was PrP^Sc^-dependent (**Fig. 4E,** lane 5). We thought that the non-convertible C;C-PrPs, e.g. those from 6C;0C to 6C;4C, cannot convert because of their unsuitable positioning of H1∼H2 for efficient refolding as discussed later, but there also was a possibility that they just cannot encounter PrP^Sc^ template in the cells. We tested the possibility by assessing their dominant negative inhibition efficiencies, as previously described [22]. Since 6C;0C to 6C;4C would not show discernible PK-res, the constructs could be used without any modification. Not unexpected, 6C;0C to 6C;5C exhibited very efficient dominant-negative inhibition on the co-expressed convertible (3F4)MoPrP (**Fig. 4F,** lanes 2-7), confirming that these C;C-PrPs do interact with template PrP^Sc^ but cannot convert into PK-res isoforms.

**FIGURE 4.**
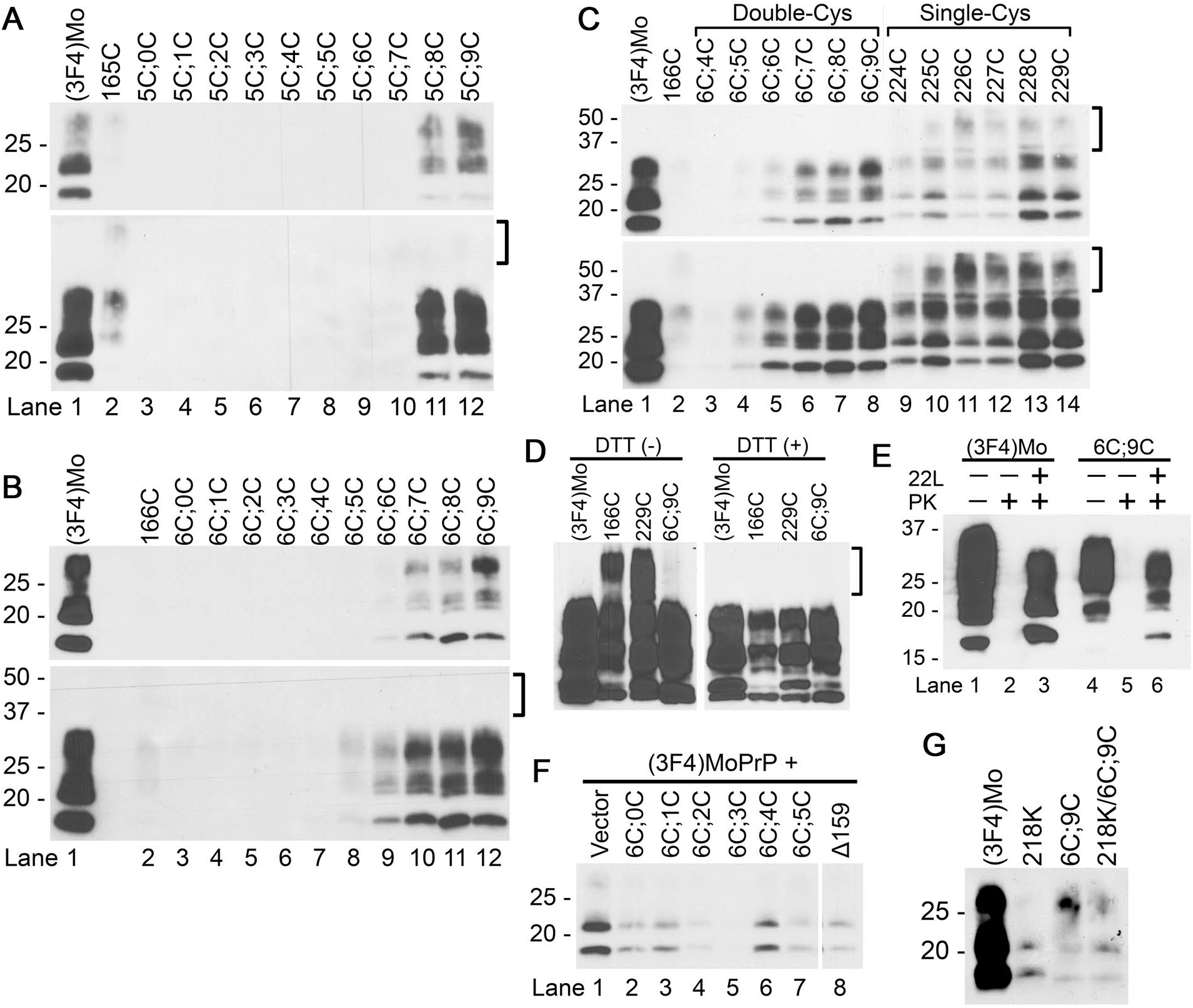
165C;C- and 166C;C-series mutants convert to PK-resistant forms (PK-res) on 22L-ScN2a. The upper and lower panels represent images of short- and long-exposure to the same PVDF membranes, respectively. The square brackets indicates the position of the dimeric form. **A and B.** PK-res of 165C;C- and 166C;C-series, respectively. Immunoblots with 3F4-mAb demonstrating levels of PK-res. Note that PK-res of C;C-series lack the dimeric forms, just as PrP^C^ isoforms. **C.** PK-res of 166C;C-series maintain the intramolecular disulfide crosslinks throughout conversion. Immunoblots with 3F4-mAb showing PK-res of 166C;C-series and mutant PrPs with a single substituted Cys, either at 166C or in Ctrm. Note that all the single-Cys constructs have substantial levels of dimeric forms, in contrast to 166C;C-series. **D.** The dimeric forms of PK-res of single-Cys mutants disappear by DTT, proving intermolecular disulfide crosslinks. Immunoblots with 3F4-mAb comparing single-Cys PrP, 166C and 229C, and a double-Cys PrP, 6C;9C. DTT (+) and (-), samples prepared with or without DTT in the sample buffer. **E**. PK-res formation of 6C;9C is PrP^Sc^-dependent. Immunoblots with 3F4-mAb demonstrating PK-res in the lysates from 22L-scrapie-infected and non-infected N2a cells, transiently-transfected with (3F4)MoPrP or C;9C. 22L + or -, samples from 22L-infected or non-infected N2a. PK + or -, samples with or without PK digestion. Note that there is no PK-res in lysates from non-infected N2a. **F**. Conversion-incompetent C;C-PrPs can interact with PrP^Sc^. Immunoblots with 3F4-mAb showing rather efficient dominant-negative inhibition effects on co-transfected (3F4)MoPrP by the conversion-incompetent C;C-PrPs, namely 6C;0-6C;5C. ?159, a deletion mutant PrP lacking the residue 159 as a control[22], was on the same membrane and unnecessary lanes were eliminated. **G**. PK-res-conversion reaction of 6C;9C is relatively resistant to inhibitory effects of Q218K substitution. Immunoblots with 3F4-mAb comparing PK-res of wild-type or 6C;9C and their Q218K counterparts. Decrements of PK-res formation by Q218K is smaller in 6C;9C than that in wild-type.

### The disulfide crosslink can suppress influences of Q218K substitution on PrP^Sc^ conversion

Lysine at the codon 219 (in human numbering; K219) is a polymorphism of human PrP well-known for protective effects against sporadic CJD [28], and the equivalent substitution in mouse PrP (Q218K) is also protective against mouse-adapted scrapie [29]. These effects were explained by the inability of K219 PrP to convert into PrP^Sc^ and its dominant-negative inhibition on the coexisting wild-type PrP [30]. As the effects of K219 was attributed to alteration in structures of the B2-H2 loop [31][32], we tested whether a mutant PrP combining 6C;9C and Q218K (218K/6C;9C) can convert to PK-res on 22L-ScN2a cells. Interestingly, 218K/6C;9C showed similar PK-res levels as 6C;9C (**Fig. 4G,** lanes 3 vs. 4), whereas Q218K PrP showed much lower levels compared to wild-type PrP (**Fig. 4G,** lanes 1 vs. 2). This suggested that the artificial disulfide crosslink of 6C;9C can suppress the effects of Q218K.

### Prediction of the existence of another C;C-PrP that can convert

The dependence of PK-res conversion of C;C-PrPs on *bona fide* PrP^Sc^ indicated that they are results of refolding reaction induced by PrP^Sc^. Tolerance of PK-res conversion reaction to specific disulfide crosslinks, namely those of 6C;5C to 6C;9C, 5C;8C and 5C;9C, seemed to be reasonably explained with a model where H1∼H2 undergoes a positional change towards Ctrm during refolding into PK-res (**Fig. 5A**): i.e. a disulfide crosslink between Cys at position 165 or 166 and Cys229 does not interfere with the refolding process (**Fig. 5B**). The model predicted the existence of another disulfide crosslink which would not interfere with the refolding reaction, bonding a more distal H1∼H2 residue and a more proximal Ctrm residue (**Fig. 5C** vs. **5D**). To test this hypothesis, we created 168C;C-PrPs (**Fig. 6A**) and assessed their conversion efficiencies in 22L-ScN2a. Expression levels of 168C;C PrPs in N2a cells were similar to that of 165C;C- and 166C;C-series mutants (**Fig. 6B**): 8C;1C showed expression comparable to (3F4)MoPrP, and 8C;4C and 8C;5C showed moderately high expression. In 22L-ScN2a cells, only 8C;4C and 8C;5C converted into PK-res forms at detectable levels (**Fig. 6C**) in a PrP^Sc^-dependent manner (**Fig. 6D**). Although we also combined 167C or 169C with Cys-scanning in Ctrm from 224 to 229 and 221 to 226, respectively, there were no discernible levels of PK-res.

**FIGURE 5.**
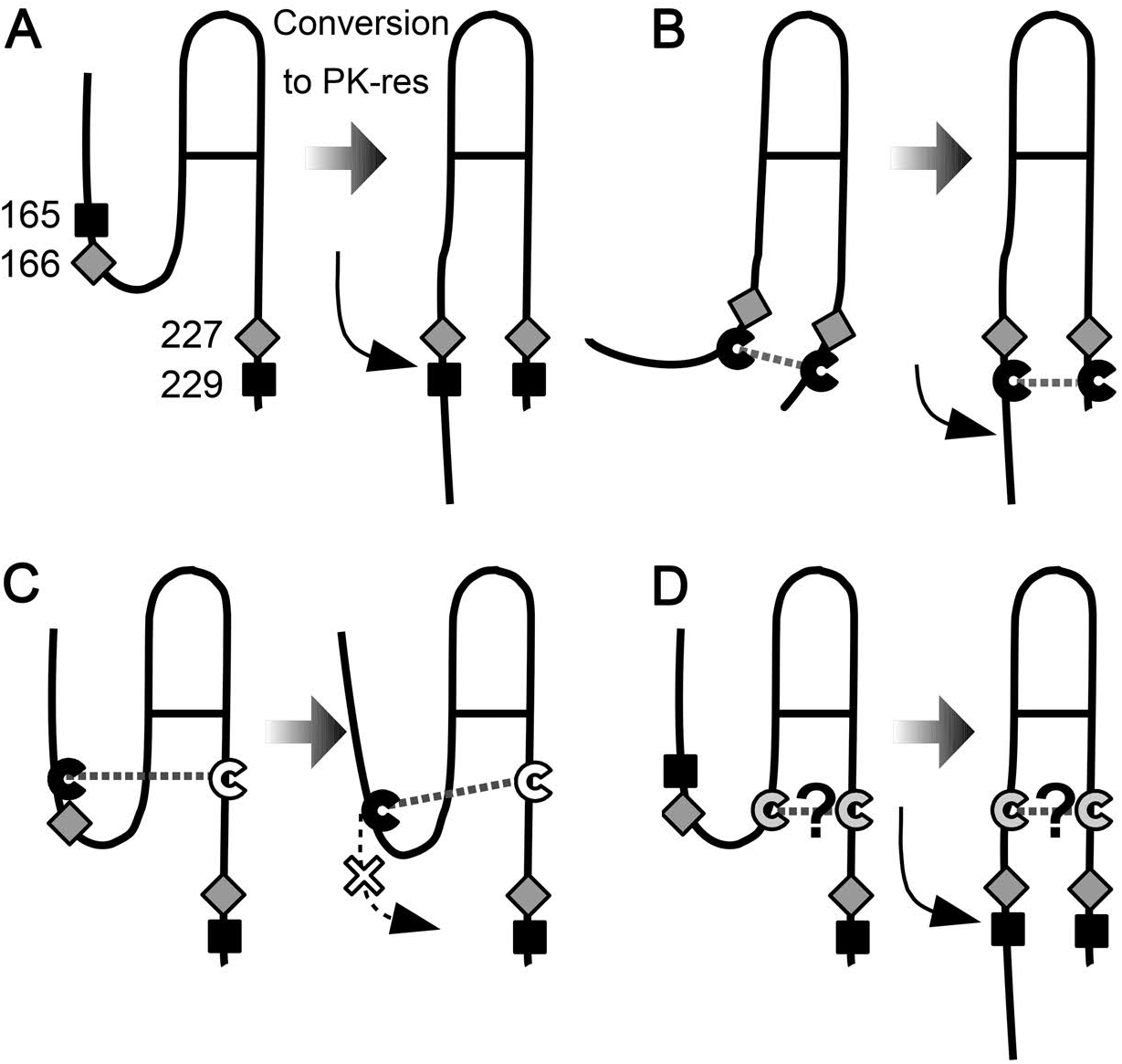
A model to explain the discrepancy between the high-expression mutants and PK-res-convertible mutants. **A.** Hypothetical positional changes of H1∼H2 in the PrP^Sc^-dependent conversion reaction. **B**. A crosslink between the residues 165 (or 166) and the distal portion of Ctrm deforms the conformation of PrP^C^ but does not severely affect the conversion because the position of H1∼H2 is suitable for conversion. **C**. A crosslink between the residues 165 (or 166) and the proximal portion of Ctrm, e.g. 220C, might inhibit PK-res conversion by hampering the positional changes of H1∼H2. **D.** Could there be a disulfide crosslink connecting the more C-terminal H1∼H2 and the more proximal Ctrm which would not hamper the conversion to PK-res?

**FIGURE 6.**
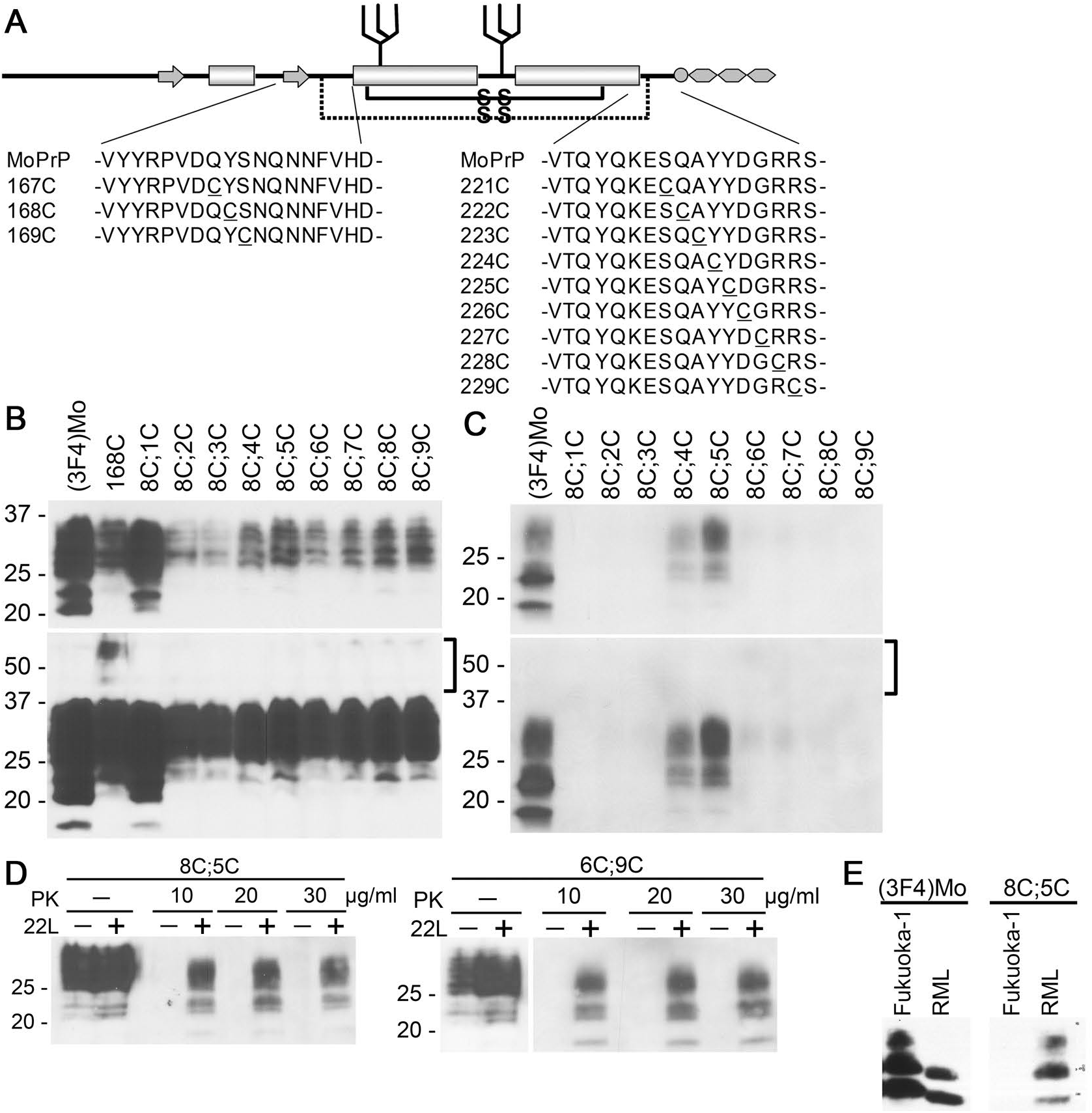
Convertible mutants of 168C;C-series supports the hypothetical positional change of H1∼H2 during conversion to PK-res. The upper panels and lower panels of blots represent images of short- and long-exposure to the same PVDF membranes, respectively. **A.** A schematic illustrating positions of substituted Cys residues of 167C;C-, 168C;C- and 169C;C-series. **B**. PrP^C^ forms of 168C;C-series showed similar banding patterns as the wild-type PrP. Immunoblots with 3F4-mAb showing expression levels and banding patterns of 168C;C-series. Square brackets, positions of dimeric forms of the mutant PrPs. **C**. Conversion capabilities of 168C;C-series. Immunoblots with 3F4-mAb demonstrating levels of PK-res of 168C;C-series. The square brackets indicates the position of the dimeric form. Like PrP^C^ forms, PK-res of 168C;C-series also lack the dimeric forms. **D**. PK-res formation of 8C;5C is PrP^Sc^-dependent like 6C;9C,. Immunoblots with 3F4-mAb comparing samples prepared from 22L-ScN2a and non-infected N2a transfected with 8C;5C or 6C;9C, digested at different concentrations of PK. Left and right panels, samples from cells transfected with 8C;5C and 6C;9C, respectively. Note that PK-res of 8C;5C is present only in 22L-ScN2a like 6C;9C. PK-digested and non-digested samples were on the same membrane and unnecessary lanes were removed. **E**. Conversion of 8C;5C into PK-res isoform is strain dependent. Immunoblots with 3F4-mAb comparing PK-res of (3F4)MoPrP and 8C;5C from transiently transfected Fukuoka1-infected or RML-infected N2a58 cells. Note that 8C;5C does not convert to PK-res in the Fukuoka1-infected cells, while it can convert in the RML-infected cells.

### Strain dependence of PK-res conversion of 8C;5C

We previously hypothesized that a PrP molecule has multiple PrP^C^-PrP^Sc^ interfaces including H1∼H2, and the usage of the interfaces are varied among different prion strains [33]. To test the hypothesis, we expressed 8C;5C on Fukuoka1- or RML-infected N2a58 cells and compared its conversion to PK-res. Surprisingly, PK-res of 8C;5C was seen only in RML-infected cells, whereas completely absent in Fukuoka1-infected cells (**Fig. 6E**).

## DISCUSSION

In this study, we have demonstrated that positional relations of H1∼H2 and Ctrm are influential on PrP^C^-PrP^Sc^ interactions and the subsequent conversion by exploiting new investigation tools, i.e. the series of systematically-designed mutant PrPs with an artificial disulfide crosslink between H1∼H2 and Ctrm. Analysis of expression levels and conversion efficiencies of C;C-PrPs in 22L-ScN2a revealed possibility of positional changes of H1∼H2 in the PrP^Sc^-guided refolding into PK-res. Besides, the positional change might be a strain-specific event, suggesting that these regions greatly contribute to the prion strain diversity. Following are the detailed discussions on the present findings:

### Expression levels of PrP^C^ isoforms of C;C-PrPs reflect their conformations

First, a variation in expression levels among C;C-PrPs, with some comparable to (3F4)MoPrP and others much less, is worthy to note. What could the variation represent? As mentioned above, a disulfide cross-link of human PrP between the residues 166 and 221 or 225 (human numbering) maintains or even stabilizes the global structure of PrP^C^ in the native conformation [25][26]. Likewise, the corresponding residues of mouse PrP^C^, residues 165 and 220, are close enough to form a stable disulfide bond [c.f. PDB ID: 2L39 [14]] and the disulfide crosslink of 5C;0C would not interfere with the native PrP^C^ conformation. This conformation possibly underlies the high expression levels of 5C;0C, because the native conformation would be thermodynamically stable with least molecular-surface hydrophobic patches which are targeted by ER or post-ER quality control systems, hence least elimination by those systems. To the contrary, the low-expression C;C-PrPs might have aberrant conformation and be actively eliminated by the quality control systems. Although the replaced residues of low-expression C;C-PrPs tend to be located too far apart to form a disulfide crosslink, possibly structural fluctuations of H1∼H2 and Ctrm allow them to crosslink and fixate PrP^C^ of C;C-PrP at an aberrant conformation.

### Implications about regional structures of PK-res

165C;C-, 166C;C- and 168C;C-series showed respective unique patterns of distribution of convertible mutants. This demonstrated that positions of the disulfide crosslinks between H1∼H2 and Ctrm are critical determinants of convertibility rather than positions of Cys itself. Since a disulfide crosslink between two regions fixates the relative positioning and local structures of the regions [34], successful introduction of artificial disulfide crosslinks to a protein without affecting the global conformation is highly informative about the regional structures of the protein. The approach was adopted in the investigation of regional structures of PrP^C^ and PrP fibrils, as well [13][25][26]. The convertible C;C-PrPs are also highly informative about the regional structures of PrP^Sc^ or PK-resistant intermediate, because positional relations of their H1∼H2 and Ctrm are compatible with the refolding process and would not be required to greatly change for the conversion. Moreover, the discrepancy between the most-highly-expressed and the most-efficiently-converted in each series (e.g. 8C;1C vs 8C;5C) is also intriguing. As discussed above, high expression levels of C;C-PrPs imply that their conformations are similar to the native PrP^C^ conformation; in other words, the positional relations of H1∼H2 and Ctrm of those mutants are suitable for the native PrP^C^ conformation. As the convertible C;C-PrPs obviously have crosslinks suitable for the conversion process, the discrepancy between the highly-expressed and the efficiently-converted is consistent with the view that a substantial positional changes of H1∼H2 toward Ctrm, from the PrP^C^-isoform position to the PrP^Sc^-isoform position, occurs in the conversion reaction in 22L-ScN2a (**Fig 6D**).

Among the convertible C;C-PrP constructs, 8C;4C and 8C;5C are particularly interesting. Kurt and colleagues reported that replacement of tyrosine at the residue 168 with aromatic residues does not affect the conversion of the mutant PrPs in *in vitro* conversion [35]. Mutant PrPs with Y168F- or Y224F-substitution also normally converts on RML-infected N2a cells [36]. Possibly aromatic-aromatic interactions between Y168 and Y224 or Y225 contribute to the conversion of wild-type PrP by bonding H1∼H2 onto Ctrm. The strain dependence of PK-res conversion of 8C;5C was the most important finding in this study, because it demonstrated strain-dependent significance of H1∼H2-Ctrm interactions for PK-res formation, which strongly supports our hypothesis that the strain diversity of PrP^Sc^ stems from varied usage of the interfaces among strains [33]. Fukuoka1 and RML PrP^Sc^ could have distinct structures in those regions. Our finding is also consistent with the strain-specific resistance of mice expressing PrP with N170S [37] regarding the strain-dependent significance of regional structures in H1∼H2. Besides, differences in the regional structures around Ctrm of PrP^Sc^ between ME7, 22L, and RML are implied by their distinct immunoreactivity [38]. The position-change model exemplifies a specific regional structure which could reasonably explain those discoveries.

The distribution of the convertible C;C-PrPs were also informative about the local structure of H1∼H2 in PrP^Sc^. As reported by Hennetin et al. [39], in the loop region of parallel β-sheet structures, i.e. “β-arches”, side chains of two successive residues often point outward of the arch, while the other regions of the β-arch have inward- and outward-facing residues alternately. Along with the proline at the residue 164, the existence of convertible mutants both in 165C;C- and 166C;C-series suggests that the residues 165 and 166 correspond to the residues at the loop region of a β-arch in PrP^Sc^, because their side chains need to point outward to form a disulfide bond with the counterpart in Ctrm. The lack of discernible PK-res in 167C;C- and 169C;C-series would be also consistent with the presence of a β-arch in the region.

Important things to be clarified through further investigation on conversion of C;C-PrPs include whether the convertible mutants can convert independently of co-existing wild-type PrP and whether they also inherit infectivity. If they can mediate infectivity and develop unique clinicopathological pictures, the C;C-PrPs might provide an insight into structure-phenotype relations of PrP^Sc^, because the local structures of H1∼H2 and Ctrm are partially predictable. In this regard, transgenic mice expressing C;C-PrPs would be even more informative.

### Effects of Q218K on the conversion efficiency of 6C;9C

K219 polymorphism of human PrP is protective against sporadic CJD but the exact underlying mechanism is yet to be identified. The partial suppression of the effects of Q218K by the disulfide crosslink of 6C;9C implied the involvement of positional relations of H1∼H2 and Ctrm. To note, although K219 of human PrP slows or modifies pathologies of sporadic CJD [28][29][31][40], new-variant CJD might not be affected or even precipitated [41][42]. This is consistent with our view that the significance of the positional relation of those regions is strain-dependent.

### Mechanism of diglycoform predominance of PrP^Sc^

Diglycoform predominance of PrP^Sc^ is characteristic of new-variant CJD and some familial CJD [12]. It also occurs in experimental transmission to elk or bank vole [20][43]. The diglycoform predominance of PK-res of C;C-PrPs in 22L-ScN2a imply that the positional relation between H1∼H2 and Ctrm of the nascent PrP^Sc^ is one determinant of the glycoform ratio. One possible mechanism is that the crosslink between H1∼H2 and Ctrm is advantageous for conversion of the diglycoform C;C-PrP. As the diglycoform of PrP^C^ isoform is much more abundant than the other glycoforms, theoretically even a small improvement in conversion efficiency of the diglycoform can change the glycoform ratio.

In conclusion, thus disulfide-crosslink scanning are unique and promising investigation tools which help locate positions of β-arches and identify their local structures. Whether β-solenoid or in-register parallel β-sheet amyloid, properties of prions can depend on those factors as long as they consist of β-strands and turns/loops[44]. Therefore, C;C-PrPs can greatly contribute to the elucidation of mysteries about prion. The intramolecular crosslink approach can be also applicable to other types of amyloids than PrP^Sc^ in theory and advance more general amyloid researches.

